# Comparison of compartmental analytical BOLD fMRI models against Monte Carlo simulations performed over cortical micro-angiograms

**DOI:** 10.1101/2024.03.06.583728

**Authors:** Jordan Charest, Mathieu Walsh, Élie Genois, Emmanuelle Sévigny, Pierre-Olivier Schwarz, Louis Gagnon, Michèle Desjardins

**Affiliations:** Department of Physics, Engineering Physics and Optics, Université Laval, Qc, Canada; Department of Physics, Université de Sherbrooke, Qc, Canada; Department of Radiology and Nuclear Medicine, Université Laval, Qc, Canada; Oncology Division, Centre de recherche du CHU de Québec - Université Laval, Qc, Canada

**Keywords:** fMRI, BOLD modeling, Monte Carlo, Two-Photon Microscopy Angiograms, Bottom-Up modelling, MRI Physics

## Abstract

BOLD fMRI arises from a physiological and physical cascade of events taking place at the level of the cortical microvasculature which constitutes a medium with complex geometry. Several analytical models of the BOLD contrast have been developed but these have not been compared directly against detailed bottom-up modeling methods. Using a 3D modeling method based on experimentally measured images of mice microvasculature and Monte Carlo simulations, we quantified the accuracy of two analytical models to predict the amplitude of the BOLD response from 1.5T to 7T, for different TE and for both gradient echo and spin echo acquisition protocols. We also showed that accounting for the tridimensional structure of the microvasculature results in more accurate prediction of the BOLD amplitude, even if the values for SO_2_ were averaged across individual vascular compartments. A secondary finding is that modeling the venous compartment as two individual compartments results in more accurate prediction of the BOLD amplitude compared to standard homogenous venous modeling, arising from the bimodal distribution of venous SO_2_ across the microvasculature in our data.

## 1. Introduction

The BOLD contrast in fMRI arises from a cascade of physiological events, including vessel size and oxygen content variations^1^. Moreover, the geometry in which MRI physics is taking place inside and outside of the cortical microvasculature^2,3^ is tortuous and anisotropic^4^. Therefore, the link between the microscopic physiological changes occurring in the cortex and the macroscopic signal measured in the MR coils is complex. Since the advent of fMRI, several biophysical models have been developed to better understand the origin of the signal and to predict its behavior with respect to different acquisition parameters and scanner magnet strengths (reviewed in Gauthier et al^5^ and Buxton et al^6^).

The Ogawa model^7^ was one of the first biophysical BOLD models developed to explain the signal amplitude, magnetic field dependence and dynamic behaviors. The balloon model^8^ was then introduced to describe some features of the hemodynamic response, and was further extended to explain other BOLD signal features (termed BOLD transients^9^) such as the initial dip^10^ and the post-stimulus undershoot^11^. Other models^12–15^ were developed with the purpose of calibrating the BOLD signal using gas challenges to compute changes in the Cerebral Metabolic Rate of Oxygen (CMRO_2_) from combined BOLD and Cerebral Blood Flow (CBF) measurements. Uludag et al ^16^ developed an analytical parametric compartmental BOLD model for both Gradient Echo (GE) and Spin Echo (SE) fMRI from 1.5T to 15T using a hybrid analytical and Monte Carlo approach to model both the intravascular and the extravascular signals. Using similar ideas, Griffeth et al ^17^ further gathered the most accepted and confirmed experimental data at that time to develop an updated compartmental model of the BOLD response for Gradient Echo at 3T. The Griffeth model was then used to validate and optimize the computation of relative changes in CMRO_2_^17^ and baseline oxygen extraction fraction (OEF)^18^ with the calibrated BOLD model. Further modeling development included consideration of the laminar aspect of the BOLD response^19–21^.

The advent of two-photon excitation microscopy^22^ has allowed researchers to measure the geometry of the cortical microvasculature^23^, as well as the partial pressure of oxygen in each vascular segment ^24^, including measurements in awake mice^25^. Using this new technology, further computational models based on Monte Carlo simulations^3,26,27^ have been developed to better account for the complex geometry and physiology of the cortical microvasculature ^28–31^. While these new modeling methods capture the complexity of the BOLD signal more accurately, their heavy computational requirements limit their utilization in statistical parametric mapping^32^ and in physiological parameter estimation, i.e. in the context of inverse problems. Therefore, simple analytical BOLD models are still broadly used in neuroimaging research.

Although the Griffeth model has been used to validate and optimize the calibrated BOLD approach to recover changes CRMO_2_ and OEF, this model was not validated in the first place and its accuracy has never been challenged against a ground-truth model. Moreover, it relies on several physical, geometrical, and physiological assumptions^1^, some of which have been proven to be more or less accurate depending on the duration of the stimulus, in particular the venous compartment blood volume changes ^33^. Therefore, in this study, our goal was to test the accuracy of the Griffeth model^17^ (and its predecessor, the Uludag model^16^), to predict the amplitude of the BOLD response at 1.5T, 3T, 4.7T and 7T as a function of the echo time (TE) in both the gradient echo (GE) and spin echo (SE) acquisition protocols. These analytical models were quantitatively compared against a Monte Carlo model computed over real microvascular angiograms of the mouse cortex. We also investigated the potential sources of error in these analytical models by comparing with hybrid versions of the Monte Carlo model.

## 2. Methods

### 2.1 Ground-truth fMRI model

We used our previous bottom-up model of BOLD fMRI based on Monte-Carlo (MC) simulations of proton diffusion in a realistic vascular network (VAN) spanning a 600 × 600 x 600 μm^3^ voxel^29^, which simulates both the intravascular and the extravascular BOLD signals. This model has been previously published and validated against experimental data and will be referred to as the full VAN model in the remainder of this paper. Extensive details about this model can be found in previous publications ^29,30,34^, and a summary is given in the following.

The realistic vascular geometry was provided by tridimensional images of the cortical vasculature measured using two-photon microscopy in 5 mice. The vascular stack was graphed^35^ and each vascular segment was labeled as artery, capillary, or vein. The diameter of each vascular segment was estimated and catalogued, as described previously^29^. For each mouse, the flow in the feeding artery was set such as to obtain a fixed perfusion of 100 ml/min/100g tissue as in our previous work ^34^, and the Cerebral Metabolic Rate of Oxygen (CMRO_2_) was computed using the method described in Gagnon et al^29^ using experimental pO_2_ measurements in the arteries and the veins for each mouse^36^. The oxygen distribution (SO_2_) in the baseline state were computed using a vascular anatomical network (VAN) model ^37,38^ together with boundary conditions obtained from the experimental pO_2_ measurements^36^.

As in a previous work in which a steady-state BOLD response was simulated^34^, we used a 10% sigmoidal arterial dilation as input to compute changes in cerebral blood flow (ΔCBF) and cerebral blood volume (ΔCBV). An intracranial pressure of 10 mmHg was assumed and the compliance parameter *β* was set to 1 for both capillaries and veins. Oxygenation changes during functional activation were then computed using the same advection code by keeping SO_2_ in the arterial inflowing segment constant and using the updated flow and volume values at each time point. CMRO_2_ was also increased using a sigmoidal temporal profile, such as to provide a ΔCBF/ ΔCMRO_2_ ratio of 3.

The magnetic field perturbation maps were computed from the tridimensional SO_2_ distribution in the vessels as described in Gagnon et al^29^ for both baseline and activated states. The resulting BOLD signal (both intra-vascular and extra-vascular) were finally computed for multiples TE’s and for both Gradient Echo (GE) and Spin Echo (SE) using the updated method described in Genois et al^30^.

### 2.2 Other BOLD models tested

In our analysis, the full VAN Monte Carlo model (labeled MC full) was considered as the ground truth against which two different analytical models were compared. Analytical models differ from our ground truth model by the fact that physiological parameters are averaged across individual vascular compartments and that the tridimensional structure of the microvasculature is not considered. In order to investigate which feature of the analytical models produced the largest error, we also implemented two hybrid versions of our Monte Carlo model in which the oxygen saturation was averaged across individual compartments but the tridimensional structure of the vasculature was considered. Therefore, a total of four models (listed below) were quantitatively compared against the ground truth model. For clarity, examples of these models are schematically presented in Fig. 1.

**Fig. 1.**
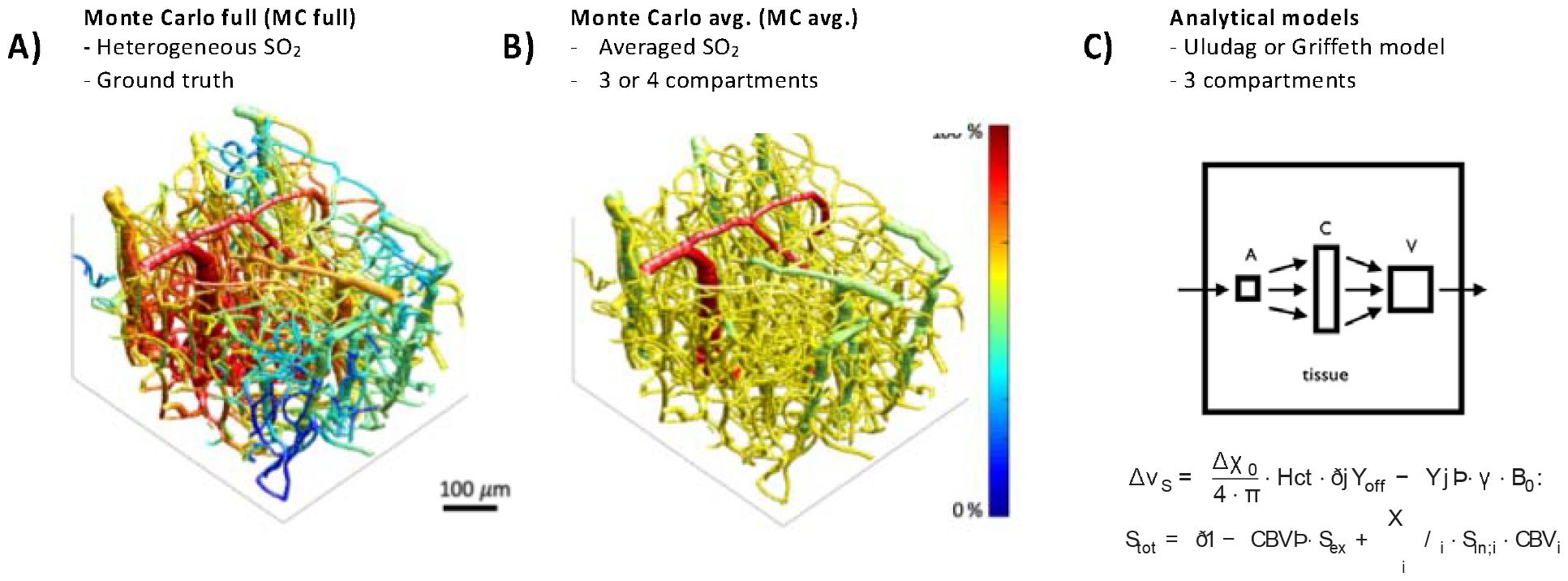
Illustration of the different simulation models used in our analysis. A) Full Vascular Anatomical Network (VAN) SO_2_ in which Monte Carlo simulations were performed. B) The mean value for SO_2_ was averaged for arteries, capillaries, and veins (3 compartments averaged) for each individual animal for both the baseline and activated states. Monte Carlo simulations of the resulting BOLD responses were performed on these 3D averaged volumes. A similar 4 compartments averaged model (not shown) was also developed but the venous compartment was further subdivided into two distinct compartments using a branching order threshold value (see Results section). C) Analytical compartmental BOLD models previously described in the literature. Two models were analyzed: the Griffeth model and the Uludag model.

1. The analytical model developed by Griffeth et al ^17^ at 3T and extended from 1.5T to 7T using physical and physiological parameters used in the literature. This model was also extended to Spin Echo BOLD modeling.
2. The analytical model developed by Uludag et al ^16^.
3. A hybrid Monte Carlo model where the real two-photon microvascular geometry was populated with mean SO_2_ values for each vascular compartment e.g. arteries, capillaries, and veins (labeled *MC avg 3 Comps*). This model was previously used in Genois et al ^30^.
4. A hybrid Monte Carlo model, similar to model 3 but where the venous compartment was further subdivided into two sub-compartments based on a vessel branching order threshold (labeled *MC avg 4 Comps*). This model was developed after a statistical analysis of the SO_2_ distribution in the venous compartment detailed in the Results section.

### 2.3 Quantitative comparison of the model

All the models were compared against the full VAN Monte Carlo model using a mean squared error (MSE) metric computed over the echo time dependence of the signal, i.e. from 0 to TE milliseconds. A schematic overview of the entire analysis framework is shown in Fig. 2.

**Fig. 2.**
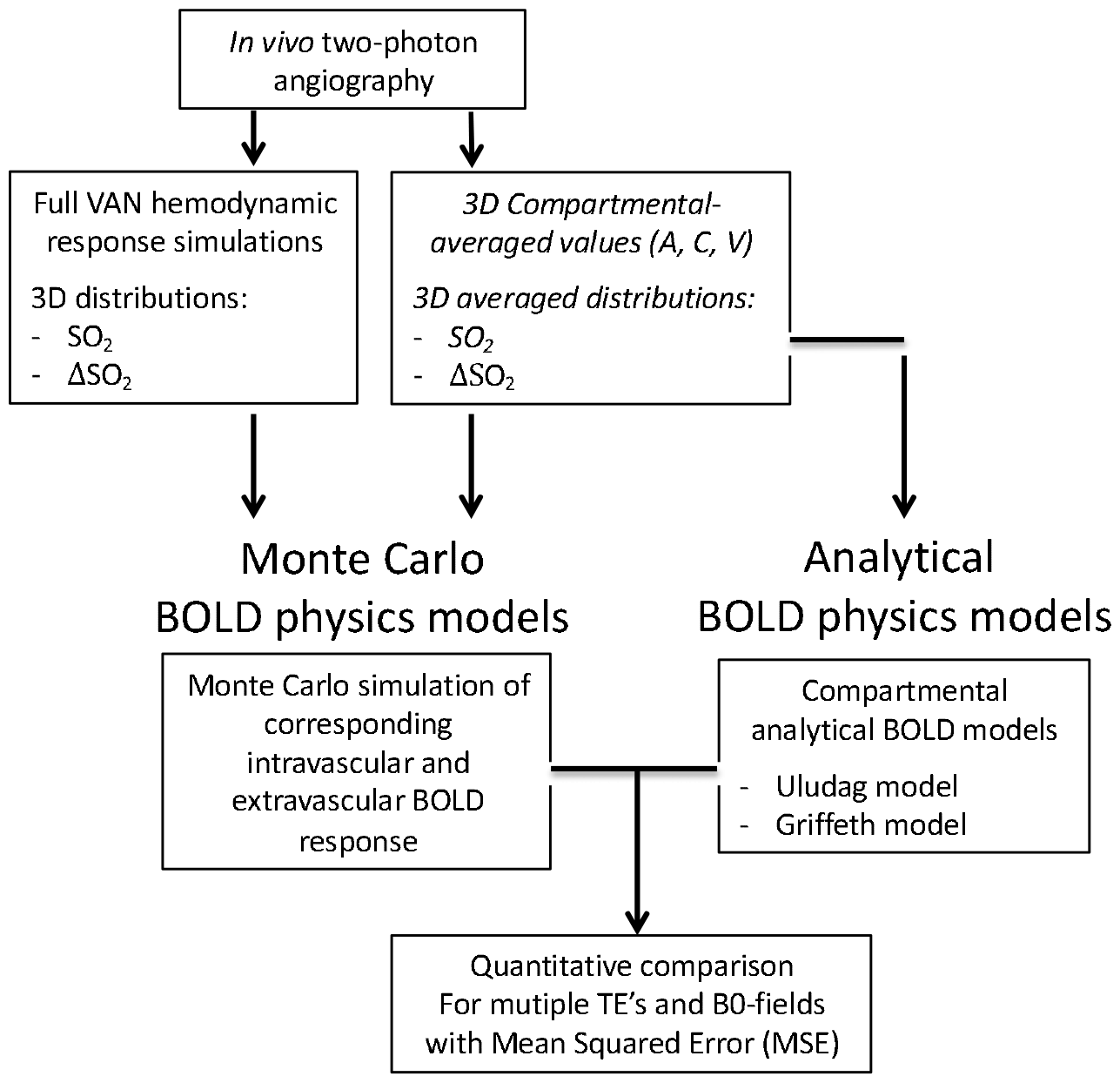
Schematic overview of the entire simulations and analysis procedure performed. The resulting BOLD response was computed for each model and was compared against the full VAN Monte Carlo model using a mean squared error (MSE) metric.

## 3. Results

### 3.1 Comparison of analytical models against Monte Carlo simulations

The echo time dependence of the BOLD signals simulated with our MC methods and the two analytical models are shown in Fig. 3A for 3T and in Fig. 3B for 7T. Traces with similar behaviors were obtained for 1.5T and 4.7T (not shown). The qualitative behavior of the echo time dependence is very consistent between the MC and analytical models, i.e. a monotonic and close to linear increasing curve is obtained in each case. The difference in the amplitudes between the analytical models and the ground truth model remains relatively small. For the case of GE at 3T (which is commonly used in fMRI experiments), the relative difference in amplitude between the analytical models and the ground truth model is about 60% (1.05% vs 0.65% BOLD amplitude at TE=30 ms). The MSE’s computed between the full VAN Monte Carlo model (considered the ground truth model in our study) and each of the analytical models are shown in Fig. 4A for GRE and Fig. 4B for SE. Results are shown for simulations at 1.5T, 3T, 4.7T and 7T.

**Fig. 3.**
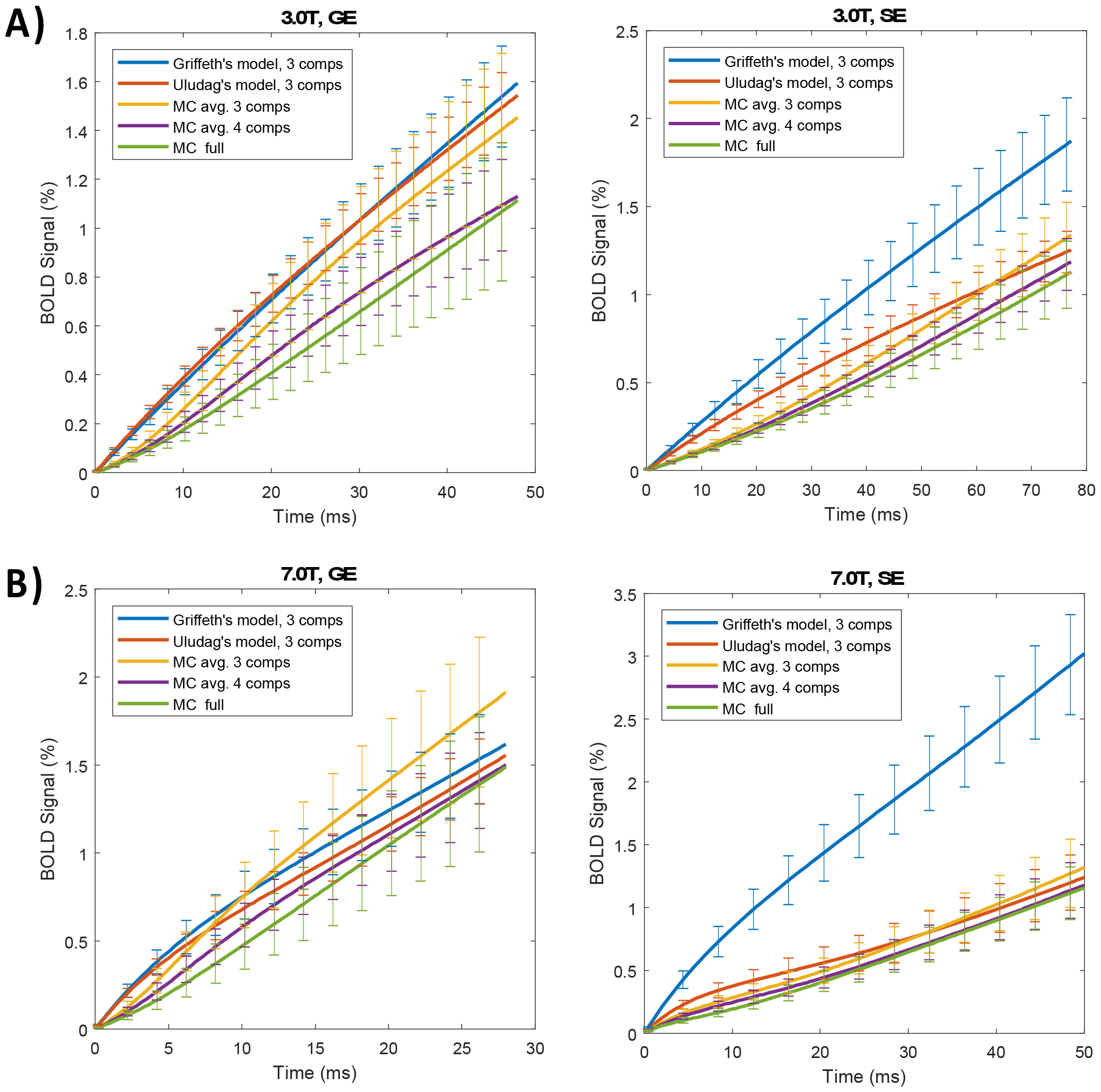
Amplitude of the BOLD response as a function of the echo time (TE) for the Griffeth analytical model, the Uludag analytical model, the 3 and 4 compartments averaged Monte Carlo models and the full VAN Monte Carlo model (each model described in Figs 1 and 2). Results are shown for both A) Gradient Echo (GE) and B) Spin Echo (SE) for B0-field values of 3T and 7T. Results represent averaged values over 5 mice and error bars represent standard error on the mean.

**Fig. 4.**
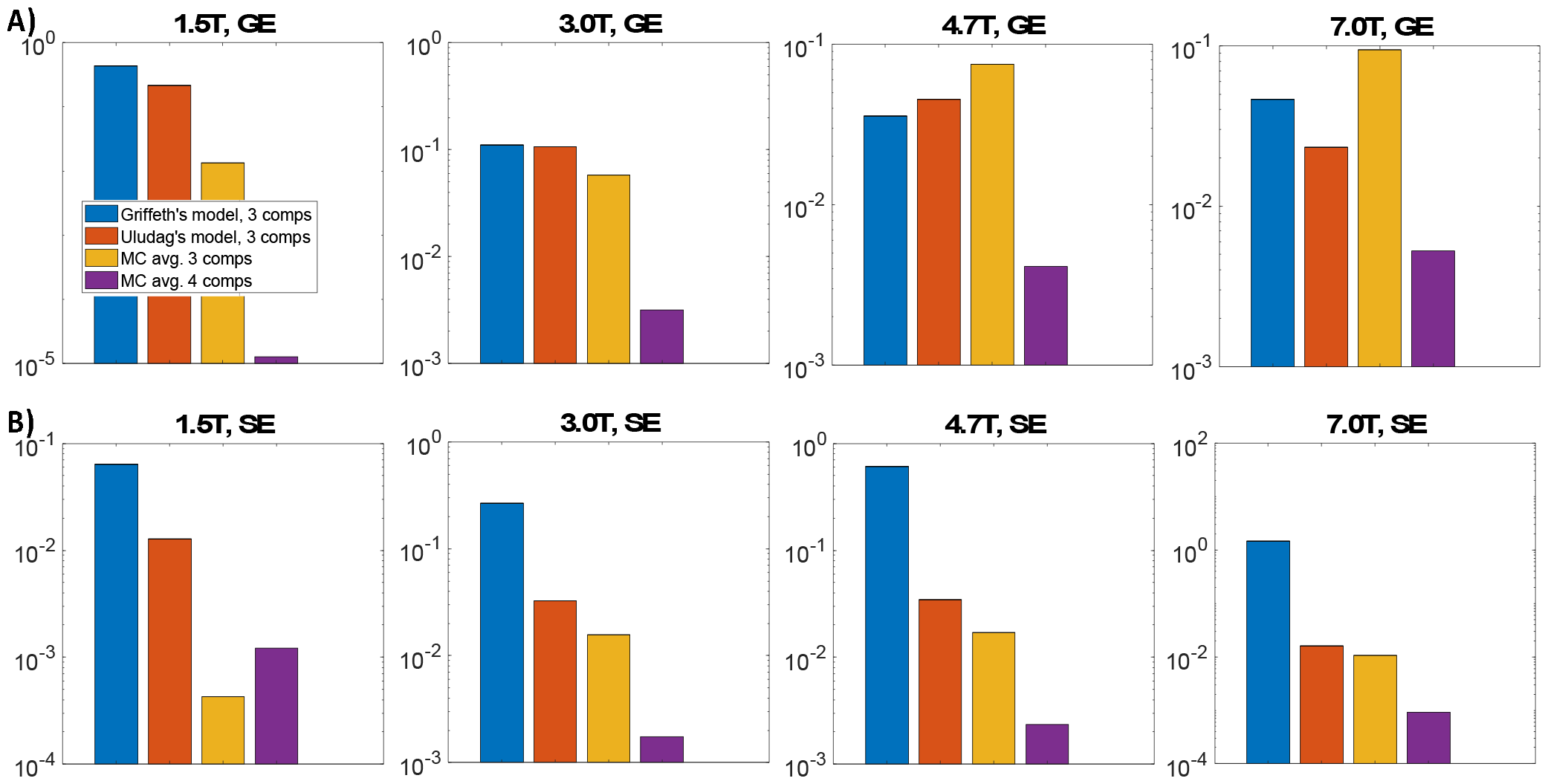
Quantitative comparison of the analytical models, the 3 compartments averaged, and the 4 compartments averaged Monte Carlo models. All models are compared against the full VAN Monte Carlo model (considered the ground truth) with a mean squared error (MSE) metric). Results are shown on a log-scale for both A) Gradient Echo (GE) and B) Spin Echo (SE) for B0-field strengths of 1.5T, 3T, 4.7T and 7T. Results represent averaged values over 5 mice.

### 3.2 Three compartments Averaged Monte Carlo simulations

To investigate the origin of the difference in amplitude between the analytical models and the full VAN Monte Carlo model, we studied a hybrid compartmental-averaged Monte Carlo model where the tridimensional structure of the vasculature was considered but was populated with mean SO_2_ values averaged across each vascular compartment (MC avg. 3 comp). Results for the echo time dependence of the BOLD signals in this case are also presented in Fig. 3. We see that for the standard GE acquisition at 3T (Fig. 3A), the relative difference in BOLD amplitude decreases from 60% (1.05% vs 0.65% BOLD amplitude at TE=30 ms) for the analytical models to 38% for the MC avg. 3 comp model (0.95% vs 0.65% at TE=30 ms). The MSE’s computed between the full VAN Monte Carlo model and the 3 compartments averaged Monte Carlo models are shown together with others models in Fig. 4A for GRE and Fig. 4B for SE. Results are shown for simulations at 1.5T, 3T, 4.7T and 7T.

### 3.3 Statistical analysis of the venous SO_2_ distributions

We investigated how to develop better analytical models by studying the origin of the difference in amplitude between the analytical models and our ground truth Monte Carlo model. The two hypothetical contributors were the loss of the tridimensional structure of the vasculature and the averaging of the physiological parameter for each vascular compartment. Our 3 compartments averaged Monte Carlo model allowed to study the effect of compartmental averaging without affecting the tridimensional structure of the vasculature. We then investigated whether considering more vascular compartments would improve our compartment averaged Monte Carlo model. As a first step, the distribution of SO_2_ across the vasculature was analysed for each individual compartment and each animal. A typical histogram of the SO_2_ distribution for the venous compartment is shown in Fig. 5A. We observe a bimodal distribution of SO_2_ with a first peak at 33 % and a second peak at 45 % for this specific animal. The bimodal appearance of the SO_2_ distribution was present in each individual animal but the exact position of the SO_2_ peaks varied from animal to animal with a mean value of 32% +/- 1% for the first peak and 42% +/- 1% for the second peak. The values for both peaks are shown in Table 1 for each animal. This bimodal behavior of SO_2_ in the venous compartment motivated us to improve the accuracy of our compartmental averaged Monte Carlo model by subdividing the venous compartment into two distinct compartments in a new 4 compartments averaged Monte Carlo model.

**Table 1.** Venous SO_2_ peaks and branching order threshold computed for each animal.

|  | **1<sup>st</sup> $\text{SO}_2$ peak (%)** | **2<sup>nd</sup> $\text{SO}_2$ peak (%)** | **Branching order Threshold** |
| --- | --- | --- | --- |
| Mouse 1 | 32 | 41 | 14 |
| Mouse 2 | 36 | 42 | 10 |
| Mouse 3 | 34 | 46 | 8 |
| Mouse 4 | 32 | 44 | 10 |
| Mouse 5 | 28 | 39 | 9 |
| Mean | 32 | 42 | 10 |
| Std Err | 1 | 1 | 1 |

**Fig. 5.**
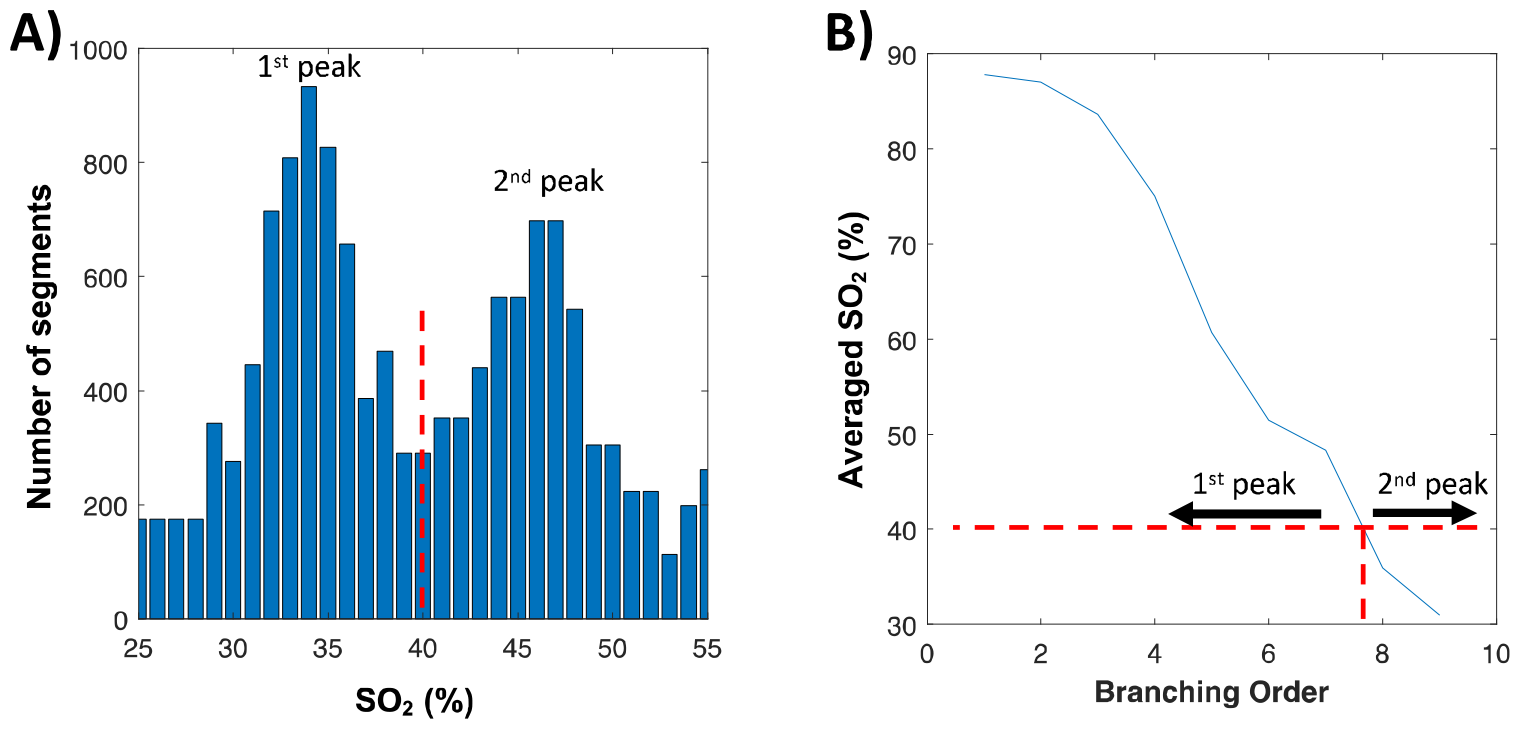
A) Typical histogram of a mouse venous SO_2_ distribution. The y-axis represents the number of vascular segments in the graph. The vertical dashed red line represents the SO_2_ threshold value used to separate the two SO_2_ peaks. B) Mean SO_2_ as a function of the branching order from the feeding artery for the same individual animal. The SO_2_ threshold identified on panel A is converted into a branching order value threshold (red dashed lines).

To subdivide the venous compartment into two distinct sub-compartments, a threshold for the SO_2_ value separating the two peaks was chosen (40% in this specific animal) and is illustrated by the red vertical line in Fig. 5A. The SO_2_ distribution as a function of the branching order from the feeding artery was plotted for each animal and is shown for the same animal in Fig. 5B. From these two plots, the SO_2_ threshold value was converted into a branching order threshold value, allowing us to classify each venous segments of the graph as part of the first or second SO_2_ peak. The branching order threshold value was 8 for this specific animal and varied from 8 to 14 (average 10 +/- 2) across all animals. The branching order threshold values are shown in Table 1 for each animal.

### 3.4 Four compartments averaged Monte Carlo simulations

Using the branching order threshold obtained for each animal venous SO_2_ distribution, the venous compartment was subdivided into two different compartments. SO_2_ values were then averaged for each of these two sub compartments, both for baseline and activated states. This subdivision allows to decrease the contribution of compartmental averaging with respect to our 3 compartments averaged Monte Carlo model. Monte Carlo simulations were performed to compute the resulting BOLD responses and were compared to the full VAN Monte Carlo BOLD response with a MSE metric. Results for the 4 compartments Monte Carlo model are shown together with results from the other models in Fig. 3 for the echo time dependence of the BOLD signal and in Fig. 4 for the MSE. We see in Fig. 3A that the relative difference in amplitude of the BOLD response drops from 38% for the MC avg. 3 comp model (0.95% vs 0.65% at TE=30 ms) to 8% for the MC avg. 4 comp model (0.70% vs 0.65% at TE=30 ms). The MSE’s obtained with the MC avg. 4 comp model are also smaller than the MSE’s obtained with the MC avg. 3 comp model for all B0-fields for GE and for B0-fields of 3T, 4.7T and 7T for SE (Fig. 4).

## 4. Discussion

### 4.1 Performance of analytical BOLD models to predict the amplitude of the BOLD response given B0-field, TE and acquisition scheme

As shown in Fig. 3, the behavior of the BOLD signal with increasing TE showed a monotonic increasing signal for each analytical and Monte Carlo model. At 3T and for GE at 7T, the differences between each model and the full VAN Monte Carlo (shown in Fig. 3) remain below 60 % with respect to the amplitude of the BOLD response. Given a steady-state hemodynamic response simulation, this result was expected. In fact, in steady state, the hemodynamic response in each vascular compartment is expected to behave relatively homogenously. Therefore, averaging the SO_2_ distributions across each individual compartment and modeling the signal with a sum of single exponentials as it is done in the Griffeth and the Uludag models does not produce large errors.

### 4.2 Accounting for the 3D structures of the microvasculature, even when SO_2_ values are averaged across individual compartments, results in more accurate BOLD response calculations

The effect of averaging the SO_2_ values across each compartment but keeping the 3D structure of the vasculature was assessed with our compartmental-averaged Monte Carlo model. We see in Fig. 4 that for both GE and SE at 1.5T and 3T, the errors computed from the compartmental averaged Monte Carlo models are lower than the ones from the analytical model. This indicates that both the compartmental averaging and the 3D structure affect the ability of the model to predict accurate BOLD responses. This result is important since it indicates that accounting for the realistic 3D structure of the cortical microvasculature, which requires much more computational cost (several hours of graphing the two-photon vascular stack and Monte Carlo proton diffusion simulations), results in more precise predictions of the BOLD response.

As shown in Fig. 4, the better performance of the compartment-averaged Monte Carlo model over the analytical models holds for both 3 compartments and 4 compartments averaging at 1.5T and 3T. The higher MSE obtained with the 3-compartment averaged MC model for GE at 4.7T and 7T can be attributed to the strong contamination of the bimodal SO_2_ distribution in the stacks that has higher effects for GE (as SE decreases venous contributions) as the B0-field increases. Accounting for this bimodal distribution with the 4-compartment averaged MC model removes this contamination and results in lower MSE values compared to the Griffeth model and the Uludag model at all B0-fields (Fig. 4). That confirms the better performance of compartmental averaged Monte Carlo models over analytical BOLD models, due to an adequate accounting of the tridimensional structure of the microvasculature.

### 4.3 Subdividing the venous compartment into two compartments improves the accuracy of the BOLD simulations

Our statistical analysis of the SO_2_ distribution performed on the individual animal level revealed a bimodal distribution of SO_2_ in the venous compartment, as shown in Fig. 5 for a specific animal. This result was consistent across all animals. As demonstrated in Fig. 4 with the MC avg. 4 Comp model, further subdividing the venous compartment into two different compartments based on a branching order threshold value resulted in lower MSE when the simulated BOLD response is compared to the full VAN Monte Carlo model. As an example, the error in BOLD amplitude for GE at 3T was about 38% for the 3 compartments averaged Monte Carlo model and dropped to 8% for the 4 compartments averaged Monte Carlo model (Fig. 3A). This result is important since it indicates that lumping the venous SO_2_ distribution in a single compartment introduce errors in the simulated BOLD response that can be removed by considering an additional vascular compartment. This result suggests that considering two individual venous compartments in an analytical BOLD model (for a total of 4 compartments) could improve its accuracy compared to a 3 compartments analytical model, without increasing the computational cost.

The bimodal distribution of SO_2_ (and therefore pO_2_) was also observed in the experimental two-photon pO_2_ measurements obtained on the same vascular stacks used in our simulations, as showed in Fig. 4B of Sakadzic et al_36_. In this same figure, the experimental SO_2_ histogram for normocapnia showed a first SO_2_ peak around 30% and a second peak around 45%, which is in good agreement with the position of the two peaks simulated with our full VAN model (32 % and 42 %) on the same vascular stacks. Additional experimental PO_2_ measurements will be required to confirm the bimodal distribution of SO_2_ in the venous compartment, but both our experimental data and our simulations suggests that this is the case and that the venous compartment must be subdivided into two separate compartments for a more accurate modeling of the BOLD response, even in steady-state. More data will also be required to determine the optimal (or minimal) number of compartments to optimize the accuracy of analytical models.

### 4.4 Comparison of the Griffeth model and the Uludag model

In addition to the model analysis performed so far, our analysis allowed us to compare the accuracy of the two analytical models (the Griffeth model and the UIudag model) to predict the BOLD response given TE, B0-field and acquisition scheme. It is important to emphasize that, while the Uludag model was designed to model the BOLD response for both GE and SE at B0-fields up to 15T, the Griffeth model was designed specifically for GE at 3T, and was naively extended to SE and to other B0-fields in the current study. As seen in Fig. 4A, in the widely used GE acquisition scheme at 3T, the MSE computed for the Griffeth model and the Uludag model (with respect to the full VAN Monte Carlo model) are not statistically different (t-test, p<0.05). We therefore conclude that both models perform equally at predicting the BOLD response amplitude at 3T for GE. Still for the GE sequence, but at B0-fields of 1.5 T and 7T, the Uludag model is shown to perform slightly better than the Griffeth model (Fig. 4A). For the SE sequence, the Uludag model outperforms the Griffeth model at all B0-fields, as expected.

### 4.5 Limitations and future studies

The development of a 4 compartments analytical model would be an interesting idea to pursue in future work, as our simulations suggested that such a model could provide a better prediction of the BOLD response amplitude without significant additional computational cost. This model could then be compared to our 4 compartments averaged MC model, and to the full VAN MC model to test its accuracy. However, the application of such an analytical 4 compartments model to predict BOLD responses in human might require more experimental data to characterize the distribution of CBV and SO_2_ in the venous compartment of the human cortical microvasculature, as differences in the venous cortical anatomy of mouse and primate are known to exist^39^.

More recently, the laminar behavior of the fMRI signal response across different cortical layers has been an active area of research and analytical multicompartment models have been developed to analyze the data and to understand the underlying contributions to the signal^19– 21,40^. Quantifying the performance of these new laminar models with our bottom-up fMRI model would be an interesting direction to explore in future work. Such a principled approach would allow to validate and potentially refine these models, in hope of better fMRI signal prediction and understanding across different cortical layers. However, this would require deeper cortical angiograms to characterize the signal across the entire cortical thickness.

This study focused on the accuracy of analytical BOLD models to predict the BOLD response in steady state, i.e. during a long continuous stimulation (>6 sec). An interesting avenue would be to analyse the accuracy of analytical models to predict the BOLD response during a short stimulus (2 sec). In this situation, the number of vascular compartments is expected to have a larger impact on the BOLD response because of the stronger effect of individual compartmental transit-time. A quantitative assessment of the accuracy of these analytical models in such conditions might allow to determine the minimal number of vascular compartments required for accurate analytical BOLD modeling during short stimulus. These validated models could then be used for Statistical Parametric Mapping in event-related fMRI, which could potentially improve the spatial localisation of cerebral activity during fMRI experiments. These validated models would ultimately provide a more quantitative and physiologically-informed measure of cerebral activity with fMRI.

## 5. Conclusion

In this work, we provided a quantitative assessment of the accuracy of analytical BOLD models to predict the amplitude of the BOLD response to a steady-state stimulus for different magnet strength (B0-field), echo time and acquisition scheme (gradient echo and spin echo). We also showed that accounting for the tridimensional structure of the microvasculature results in more accurate prediction of the BOLD amplitude, even if the values for SO_2_ were averaged across individual vascular compartments. In addition, we found that subdividing the venous compartment into two individual compartments results in better prediction of the BOLD amplitude, arising from the bimodal distribution of venous SO_2_ across the microvasculature in our data.

### Appendix

Both analytical models (Griffeth and Uludag) are used to compute the dBOLD signal given by:

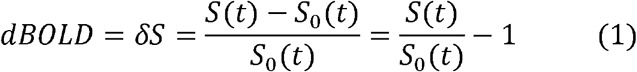

where *S*(*t*) is the signal during activation at echo time t=TE and *S*_0_(*t*) is the signal at baseline. The signal for a given compartment *i* is given by:

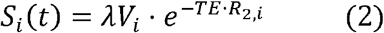

Where *λ* = 0.87/0.89 is the relative proton spin density between intravascular and extravascular tissue^41^. The total signal is the sum of the signal of every compartment, with averaged *SO*_2_ values across each compartment as previously detailed:

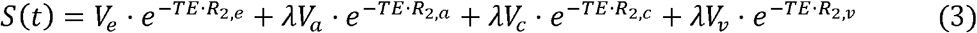

#### Uludag et al.^16^

Taking the above equations, the relaxation rate is split between the intrinsic relaxation rate 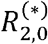 and the additional contributions to relaxation caused by deoxyhemoglobin in the vessels:

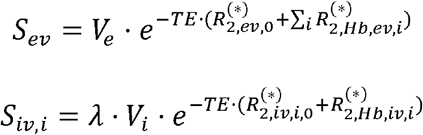

These equations hold for signals both during baseline and activation. The following equations give the intrinsic relaxation rates:

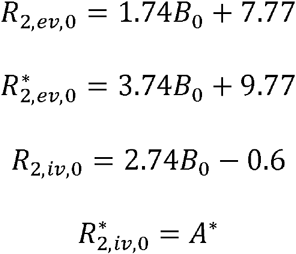

The contributions to the intravascular signal from deoxy-Hb for compartment *i* are given by:

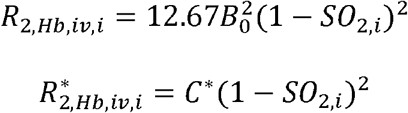

Both for baseline and activation. Both *A*^*^ and *C*^*^ are constants obtained from fits on experimental data^29-30^ (more information below). For the extravascular signal, the contributions are given by a polynomial fit over results of Monte-Carlo simulations^16^:

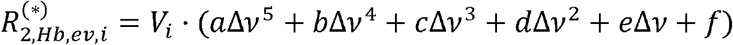

Where Δ𝒱 is the Larmor frequency shift induced by blood susceptibility:

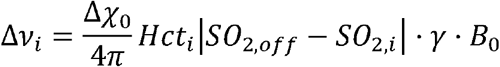

Δ𝒳_0_ = 4*π* · 0.264 ppm is the susceptibility of deoxygenated blood, Hct is the hematocrit for each compartment (0.44 for arteries and veins and 0.33 for capillaries^42^), *SO*_2,*off*_ = 0.95 is the oxygen saturation that produces no susceptibility difference between intravascular and extravascular tissue, and γ = 2*π* · 42.6 MHz/T is the gyromagnetic ratio of protons.

#### Griffeth et al.^17^

The model developed by Griffeth et al. also uses equations 1-3 to compute the dBOLD signal. Taking equation 1 and expanding it, we obtain:

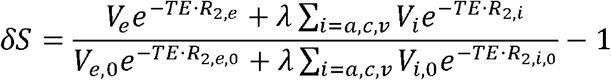

We multiply the fraction by 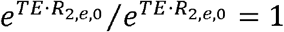

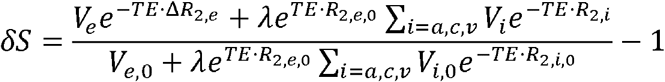

We multiply the numerator sum by 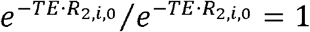

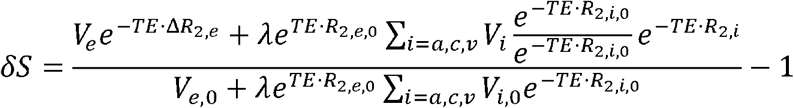

Griffeth et al. defines the quantity *ε* as the ratio between intravascular and extravascular baseline signals:

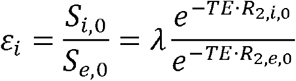

This quantity can be used to simplify the previous equation:

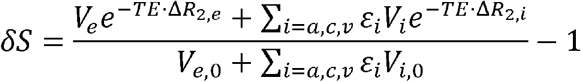

Values for the relaxations rates 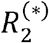 are needed to compute the signal, with the relaxation rates for GRE denoted by 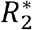 and those for SE denoted by *R*_2_. For the intravascular compartments (a, c, v) Griffeth et al. uses a quadratic model for 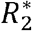 dependent on oxygen saturation *SO*_2_ :

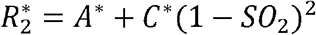

The dependence on hematocrit of *SO*_2_ used in Griffeth et al. was neglected in favor of values for *A*^*^ and *C*^*^ taken from the literature up to 7.0T^29-30^. Values above 7.0 T are obtained from a linear extrapolation.

The values for *SO*_2_ and volume, during activation and baseline for each compartment, are determined using our own advection code. The same values are used for the Monte-Carlo simulations as well as for the VAN and analytical models. The change in relaxation rate can thus be computed using the values of *SO*_2_ for each compartment.

For the extravascular compartment, the change in relaxation rate is obtained from a linear combination of contributions from the intravascular compartments.

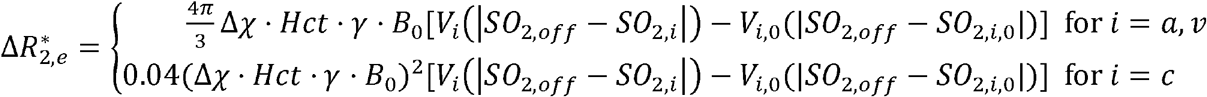

And the total change in relaxation rate is the sum of the three contributions. The baseline 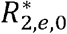 value is taken from the literature^43^.

Griffeth et al.’s model is specifically tailored for GE. As such, the following equations were borrowed from Uludag et al.’s model to make SE computations possible:

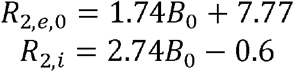

With the rest of the model remaining as it is.

## Acknowledgements

We thank David Boas, Anna Devor and Sava Sakadzic for sharing their microscopy data. This work was supported by an NSERC Discovery Grant (RGPIN-2018-06138) and by a FRQS J1 Grant (309938).

## Data and code availability statement

The presented data and code developed for this study are openly available for sharing upon request by emailing the Corresponding Author.

## Credit authorship contribution statement

J. Charest, M. Walsh, E. Genois and P-O. Schwarz : Performed simulations and analyzed results. L. Gagnon: Conceptualization, methodology, performed simulations, analyzed results, supervised the project and wrote the manuscript. M. Desjardins: Conceptualization, methodology, supervised the project and wrote the manuscript. E. Sévigny: Wrote the manuscript.

